# Default and Control networks connectivity dynamics track the stream of affect at multiple timescales

**DOI:** 10.1101/2020.06.06.137851

**Authors:** Giada Lettieri, Giacomo Handjaras, Francesca Setti, Elisa Morgana Cappello, Valentina Bruno, Matteo Diano, Andrea Leo, Emiliano Ricciardi, Pietro Pietrini, Luca Cecchetti

**Affiliations:** Social and Affective Neuroscience Group, IMT School for Advanced Studies Lucca, Lucca, Italy; Molecular Mind Laboratory, IMT School for Advanced Studies Lucca, Lucca, Italy; MANIBUS Lab, Psychology Department, University of Turin, Turin, Italy; Department of Psychology, University of Turin, Turin, Italy; Department of Translational Research and Advanced Technologies in Medicine and Surgery, University of Pisa, Pisa, Italy

**Keywords:** default mode network, control network, naturalistic stimulation, fMRI, affect

## Abstract

In everyday life the stream of affect results from the interaction between past experiences, expectations, and the unfolding of events. How the brain represents the relationship between time and affect has been hardly explored, as it requires modeling the complexity of everyday life in the laboratory setting. Movies condense into hours a multitude of emotional responses, synchronized across subjects and characterized by temporal dynamics alike real-world experiences.

Here, we use time-varying intersubject brain synchronization and real-time behavioral reports to test whether connectivity dynamics track changes in affect during movie watching. Results show that polarity and intensity of experiences relate to connectivity of the default mode and control networks and converge in the right temporo-parietal cortex. We validate these results in two experiments including four independent samples, two movies, and alternative analysis workflows. Lastly, we reveal chronotopic connectivity maps within temporo-parietal and prefrontal cortex, where adjacent areas preferentially encode affect at specific timescales.

## Introduction

Emotions are intense and immediate reactions of the body and the brain to an external or internal event occurring in the present, happened in the past, or may occur in the future. In daily life, the temporal trajectory of emotions is characterized by regularities in features, such as the time to rise, the duration, or the probability of resurgence (Kuppens and Verduyn, 2017), which inform mental models of emotion transitions and support affective forecasting (Thornton and Tamir, 2017). Whence, it appears that the study of diachronicity is crucial for building a more comprehensive understanding of emotion processing and conceptualization in humans. Indeed, over the last years, increasing attention has been dedicated to the study of the temporal characteristics of affect using behavioral and experience sampling methods (Kuppens et al., 2010; Trampe et al., 2015; Verduyn et al., 2015), allowing researchers to track emotions over days, weeks, or months. In sharp contrast, studies on the brain correlates of affective dynamics are still limited (Waugh and Schirillo, 2012; Costa et al., 2014; Verduyn et al., 2015; Résibois et al., 2017; 2018) and have employed static or relatively brief stimuli (Posner et al., 2009; Baucom et al., 2012; Kim et al., 2017) that may not be adequate to account for the complex temporal trajectory of lifelike experiences (Waugh and Schirillo, 2012).

A possible strategy to overcome this limitation is measuring brain activity elicited by movies using functional magnetic resonance imaging (fMRI). Indeed, movies are ecological and dynamic stimuli that synchronize brain response across individuals (Hasson et al., 2004) and elicit a wide variety of emotional states in a few hours (Schaefer et al., 2010). In addition, movies represent an effective tool for tracking time-varying brain connectivity (Nastase et al., 2019) and revealing the interplay among large-scale networks. Promising evidence in this regard comes from a work that, using relatively brief movie excerpts (∼8 minutes), demonstrated the association between connectivity of salience and amygdala-based networks and the perceived intensity of sadness, fear, and anger (Raz et al., 2016).

Also, brain connectivity dynamics well exemplify the tight relationship between emotion and cognition, as the same brain region can interact with distinct functional networks over time, and the same network can support multiple mental processes (Pessoa, 2017; 2018). In line with this, studies combining naturalistic stimulation with brain connectivity measurements show that the default mode network (DMN) contributes to information encoding (Simony et al., 2016), memory representations (Chen et al., 2017), and prediction of future events (Antony et al., 2021). Interestingly, it has been argued that the same network is also crucial for constructing emotional experiences (Satpute and Lindquist, 2019) and integrating the self with the social world (Yeshurun et al., 2021).

Here, we test the hypothesis that changes in DMN connectivity would reflect the temporal properties of emotion dynamics experienced by individuals. In a first experiment, we explore whether, during the viewing of a 2-hour movie (Forrest Gump; Hanke et al., 2016), subjective reports of the affective experience explain brain connectivity dynamics in independent participants watching the same film. To rule out the possibility that results depend on the selected stimulus, we validate our findings in a second experiment by acquiring emotional reports and brain data in other individuals presented with a different live-action movie (i.e., 101 Dalmatians). Also, given the impact of analytical choices on brain imaging results (Botvinik-Nezer et al., 2020), we test the generalizability of findings by preprocessing fMRI data using two alternative workflows. Lastly, in light of topographies revealed in high-order associative areas (Baldassano et al., 2017), we test the existence of a chronotopic organization (Protopapa et al., 2019) in regions encoding the stream of affect.

## Materials and Methods

### Experiment 1: Behavioral Study

Behavioral data have been previously reported (Lettieri et al., 2019) and made freely available at https://github.com/psychoinformatics-de/studyforrest-data-perceivedemotions. In brief, twelve Italian native speakers (5F; mean age 26.6 years, range 24–34) provided ratings of their moment-by-moment emotional experience while watching the Forrest Gump movie (R. Zemeckis, Paramount Pictures, 1994; for details see Supplementary Data).

They were presented with an Italian dubbed and edited version of Forrest Gump, split into eight movie segments ranging from 11 to 18 minutes. Subjects were asked to report (i.e., 10Hz sampling rate) the type and intensity of their inner emotional state using six emotion categories (happiness, surprise, fear, sadness, anger, and disgust; Ekman, 1992). As the complexity of the experience elicited by Forrest Gump cannot be merely reduced to six emotions, we allowed participants to report combinations of categories throughout the movie, simultaneously. Our approach revealed that behavioral ratings captured 15 different affective states (see Lettieri et al., 2019). Starting from categorical descriptions, we then obtained the underlying affective dimensions - i.e., polarity, complexity, and intensity - using principal components analysis (see Supplementary Data).

For each subject, the behavioral data acquisition’s overall duration was 120 minutes (72,000 timepoints). Stimulus presentation and recording of the responses were implemented in Matlab (R2016b; MathWorks Inc., Natick, MA, USA) and Psychtoolbox v3.0.14 (Kleiner et al., 2007).

### Experiment 1: fMRI Study

Brain activity elicited by the watching of Forrest Gump was obtained from the studyforrest project - phase II (n=14, German native speakers; Hanke et al., 2016). Details on brain data acquisition and preprocessing can be found in Supplementary Data.

To track time-varying brain connectivity, we opted for the intersubject functional correlation (ISFC) approach, which computes the correlation between signal change of a specific brain region in one subject with the activity of all other regions in other subjects (Simony et al., 2016). Compared to other functional connectivity measures, ISFC demonstrates a higher signal-to-noise ratio as it inherently filters out activity not related to stimulus processing and artifacts not correlated across brains. In addition, ISFC can be used to study dynamic reconfigurations of brain networks using a moving-window approach. We estimated the static and time-varying ISFC (tISFC) associated with the watching of Forrest Gump using custom MATLAB scripts, freely available at https://osf.io/v8r9w/?view_only=6a8ac3e3c6ba43d0a205146b4975ebb4.

As a first step, we selected the atlas provided by Schaefer and colleagues (2018) and parcellated each subject’s brain into 1,000 cortical regions of interest (ROIs). We opted for this atlas as, compared to other parcellations (e.g., Shen et al., 2013), it provides a finer-grained segmentation of the cerebral cortex that, at the same time, reflects the seven resting state networks (Yeo et al., 2011): visual, somato-motor, ventral attention, dorsal attention, limbic, control, and default mode network. In addition, as Schaefer parcellation does not include subcortical regions, we used Harvard-Oxford subcortical atlas (Makris et al., 2006) to define 14 additional ROIs (7 per hemisphere): Nucleus Accumbens, Amygdala, Caudate, Hippocampus, Pallidum, Putamen, and Thalamus. Thus, in total, the parcellation we adopted comprises 1,014 areas.

To compute static ISFC, the time course of brain activity was extracted from each ROI of one subject and correlated using Pearson’s coefficient with the remaining 1,013 regions of all other subjects. By repeating this procedure for each of the 14 participants, we generated 91 (i.e., number of all possible combinations of subjects) correlation matrices. We then applied Fisher z-transformation and averaged correlation values across subjects to produce a group-level ISFC map.

To estimate tISFC, one has first to determine the width of the moving-window and the degree of overlap between adjacent windows. Both parameters are often chosen arbitrarily, even though they may significantly affect the results. To limit arbitrariness, we estimated the optimal window width as a function of the intersubject correlation in affective dimensions obtained from the behavioral experiment. Specifically, we computed the average intersubject correlation of affective dimension timeseries for all the possible window widths, ranging from (1) 20 to 250, (2) 20 to 500, (3) 20 to 750, and (4) 20 to 1,000 timepoints (i.e., from 40s to ∼33m) and having 33% of overlap (Figure 1A and 1B). Of note, although the upper bound of the explored intervals (i.e., 250, 500, 750, 1,000) is arbitrarily selected, the optimal window size is determined by a data-driven approach. Specifically, the optimal timescale was defined for each range as the point at which the intersubject correlation curve starts to flatten (i.e., knee point); namely the point representing a reasonable trade-off between the agreement across participants and the window width (Figure 1B). Depending on the dynamics of emotional ratings, this procedure provides up to four optimal window widths (i.e., one for each range), and it was repeated for each affective dimension separately. The reason for exploring four timescales is that distinct brain regions segment and process narratives at multiple temporal resolutions (Baldassano et al., 2017), and a similar mechanism could be hypothesized for the processing of affective information.

**Figure 1.**
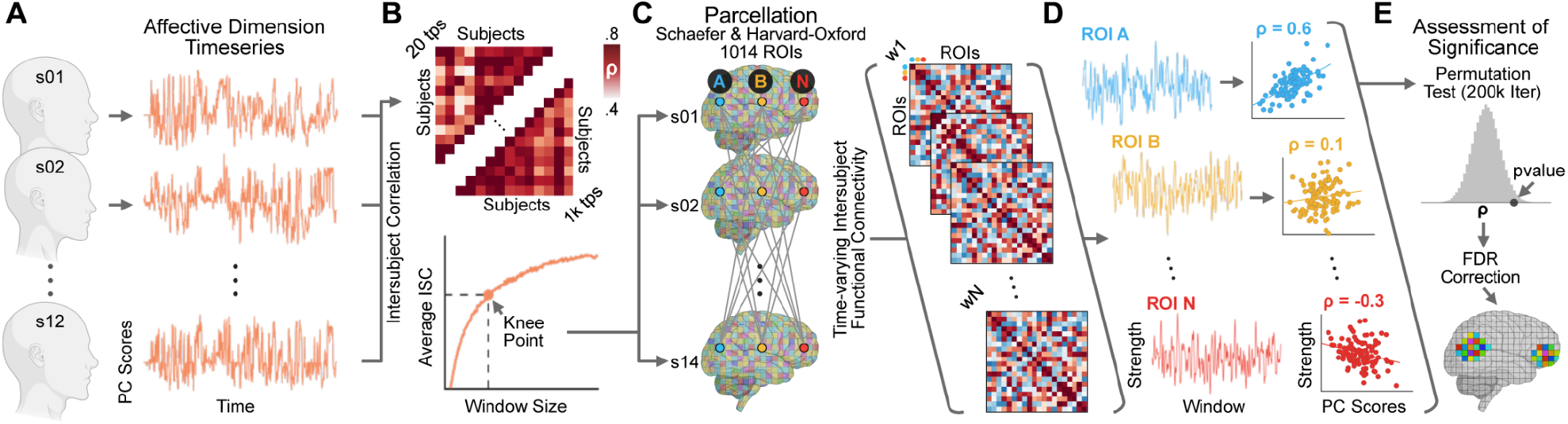
summarizes how we compute the association between affective dimensions extracted from behavioral ratings and time-varying connectivity strength of 1,014 brain regions. Each subject provided ratings of the affective experience during movie watching. We then applied principal component analysis to reveal single-subject affective dimension timeseries (polarity, complexity and intensity; panel **A**). Affective dimensions were correlated across subjects using a sliding window approach with multiple window sizes, ranging from 20 to 1,000 timepoints. Based on the tradeoff between window size and average intersubject correlation in behavioral ratings we selected four optimal width (i.e., knee point; panel **B**) for the subsequent estimation of brain connectivity dynamics. Brain activity was extracted from 1,014 regions of interest and time-varying intersubject functional correlation (tISFC) was computed following Simony et al., 2016 (panel **C**). For each brain area we estimated changes in connectivity strength on the complete graph. We correlated (Spearman’s ρ) the obtained timeseries with the timecourse of group-level affective dimensions (panel **D**). The assessment of significance was performed using a non-parametric permutation test and false discovery rate correction (Benjamini and Yekutieli, 2001) for multiple comparisons was then applied (panel **E**).

Once optimal window widths were established, we computed the tISFC as the Pearson’s correlation between each ROI of one subject and all the other ROIs of the other subjects in each window (Figure 1C). We then applied Fisher z-transformation and averaged correlation values across subjects to produce a group-level timeseries of brain connectivity (i.e., tISFC) associated with movie watching.

The primary aim of our study was to relate changes in tISFC to variations in the perceived affective state during movie watching. Therefore, for each ROI we computed connectivity strength as the sum of the Z-transformed correlation values at each timepoint (Figure 1D). We thus obtained 1,014 timeseries expressing the relationship of each ROI with the rest of the brain throughout the movie. These timeseries were then correlated using Spearman’s ρ coefficient with the group-level timecourse of affective dimensions, after those were downsampled to the optimal window widths using a moving-average procedure. To assess the significance of the association (Figure 1E), we used a non-parametric permutation test based on timepoint shuffling of affective dimensions (200,000 iterations; minimum two-tailed p-value: 1.0e-5). At each iteration, randomized affective dimensions were correlated with ROIs connectivity strength, providing null distributions of Spearman’s correlation values. The position of the actual association in the null distribution determined the two-tails level of significance. To correct for multiple comparisons, we pooled together the 1,014 p-values obtained from each affective dimension and timescale (12 in total; 4 for polarity, 4 for complexity and 4 for intensity) and used the false discovery rate algorithm under dependency assumptions (Benjamini and Yekutieli, 2001) with a threshold of 0.05.

In addition, the selected parcellation allowed us to characterize the relationship between changes in affective dimensions and functional brain networks. To graphically represent the involvement of each network, we first used classical Multi-Dimensional Scaling (MDS; Torgerson, 1952) on the group-level ISFC to map functional distances between the 1,014 brain regions during movie watching. In the MDS plot, each region was color-coded per network membership and scaled in size and transparency depending on the log p-value of the association between connectivity strength and changes in affective dimensions. Results of this procedure are represented in Figure 2.

**Figure 2.**
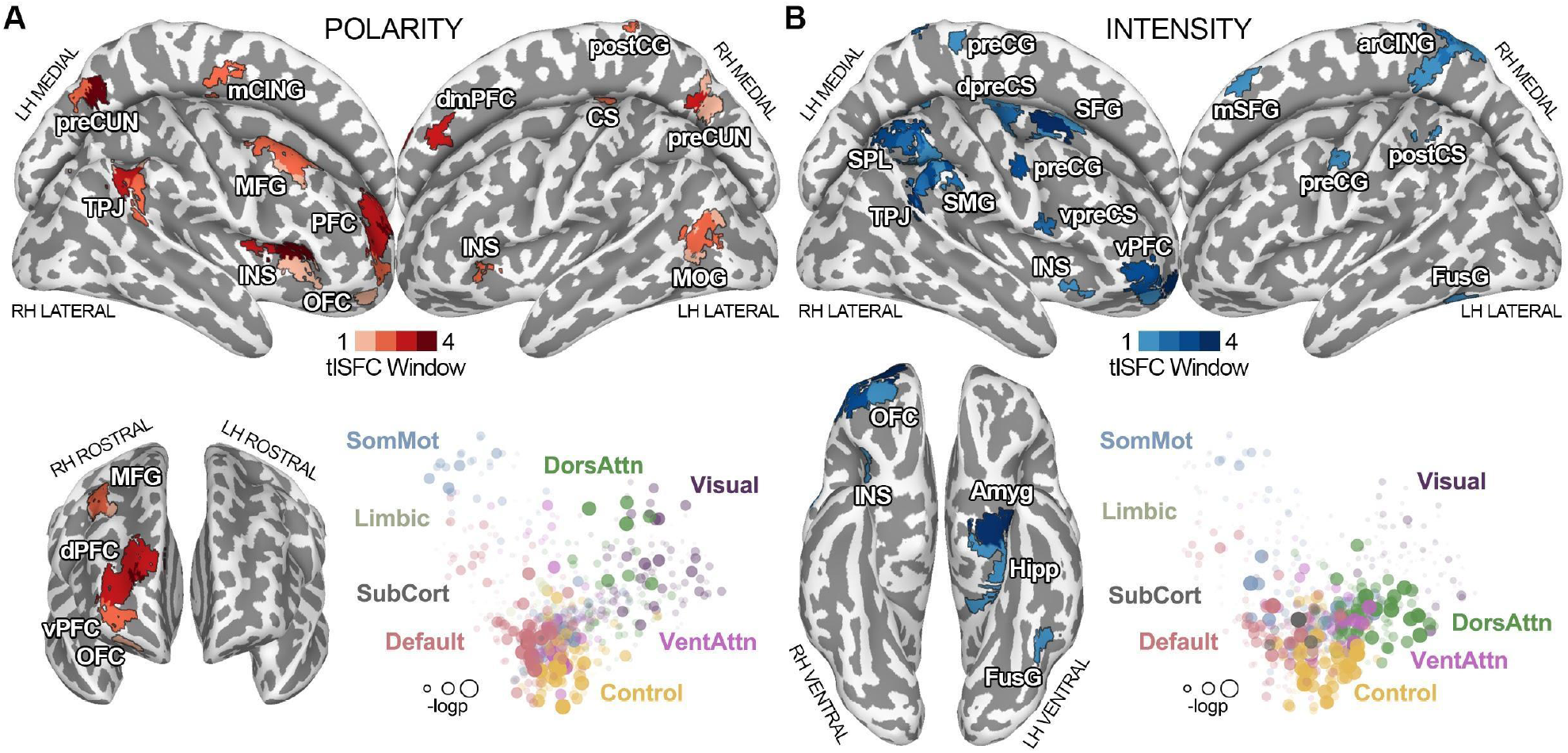
Panel **A** shows results for the connectivity strength of brain regions associated with changes in the polarity of the affective experience at all explored timescales. Grading of the color red indicates the number of window widths in which the connectivity strength of a region was associated with polarity. In the multidimensional scaling (MDS) plot, located in the lower part of the figure, each brain area (i.e., dot) is color-coded depending on network membership, whereas its size and transparency are scaled according to the -log(pvalue) of the relationship with polarity. In panel **B**, connectivity strength of brain regions associated with changes in the intensity of the affective experience at all explored timescales is reported. Grading of the color blue indicates the number of window widths in which the connectivity strength of a region was associated with intensity. In the MDS plot, each brain area (i.e., dot) is color-coded depending on network membership, whereas its size and transparency are scaled according to the -log(pvalue) of the relationship with intensity. LH = left hemisphere; RH = right hemisphere; CS = central sulcus; arCING = ascending ramus of the right cingulate sulcus; Amyg = amygdala; Hipp = hippocampus; preCUN = precuneus; TPJ = temporoparietal junction; INS = insula; dPFC = dorsal prefrontal cortex; vPFC = ventral prefrontal cortex; mCING = mid cingulate cortex; FusG = fusiform gyrus; MFG = middle frontal gyrus; dmPFC = dorsomedial prefrontal cortex; MOG = middle occipital gyrus; PFC = prefrontal cortex; preCG = precentral gyrus; postCG = postcentral gyrus; postCS = postcentral sulcus; dpreCS = dorsal precentral sulcus; vpreCS = ventral precentral sulcus; SFG = superior frontal gyrus; mSFG = medial superior frontal gyrus; SPL = superior parietal lobule; SMG = supramarginal gyrus; OFC = orbitofrontal cortex; DorsAttn = dorsal attention network; VentAttn = ventral attention network; SomMot = somatomotor network; SubCort = subcortical network.

Lastly, to test the existence of a chronotopic organization of affect associated with connectivity dynamics, we exploited the multiple timescales derived from the optimal window estimate. In brief, we first estimated the significance of the associations between connectivity and affective dimensions at all explored timescales, as described above. A logical OR across timescales was then computed for each affective dimension and a winner-takes-it-all criterion on the log-transformed p-value maps established the preferred timescale of each brain region (Figure 4).

### Experiment 2: Behavioral Study

Twenty-one Italian native speakers (13F; mean age 29.4 years, range 23–44) provided ratings of their emotional experience while watching, for the first time, an edited version of the live-action movie 101 Dalmatians (S. Herek, Walt Disney Pictures, 1996). All participants were clinically healthy and had no history of any neurological or psychiatric condition. Each subject was informed about the nature of the research and gave written informed consent to participate in the study. The local Ethical Review Board approved the experimental protocol and procedures (CEAVNO: Comitato Etico Area Vasta Nord Ovest; Protocol No. 1485/2017) and the research was conducted in accordance with the Declaration of Helsinki.

Differently from experiment 1, in the validation dataset, participants were asked to report moment-by-moment (10Hz sampling rate) the pleasantness or unpleasantness and intensity of the experience on a continuous scale ranging from -100 (extremely negative) to +100 (extremely positive). For each subject, raw ratings were interpreted as a measure of polarity, whereas its absolute value was considered a measure of intensity. For stimulus preparation and task implementation, please see Supplementary Data.

### Experiment 2: fMRI Study

Brain activity elicited by the watching of 101 Dalmatians was obtained from ten healthy Italian subjects (8F; 35±13 years), instructed to simply inhibit any movement and enjoy the movie. Individuals participating in the MRI study did not take part in the behavioral experiment. Also, they watched the movie 101 Dalmatians for the first time inside the scanner. The fMRI acquisition lasted 1 hour and was split in six runs (3T Philips Ingenia scanner; GRE-EPI: TR=2000ms, TE=30ms, FA=75°, 3 mm isotropic voxel). The Ethical Committee of the University of Turin (Protocol No. 195874/2019) approved the fMRI study.

Considering the high variability in results related to analytical choices (Botvinik-Nezer et al., 2020), we used two alternative pre-processing pipelines to analyze the validation dataset (i.e., workflow A and B), which are detailed in Supplementary Data.

To evaluate the reliability and generalizability of the association between tISFC and the timecourse of affective dimensions reported in experiment 1, we tested whether brain connectivity dynamics explain changes in polarity and intensity in 101 Dalmatians. To this aim, we followed the procedure detailed in *Experiment 1 - fMRI study* (see also Supplementary Data) and used a conjunction analysis of results obtained from experiment 1 to restrict the search for significance. Specifically, we selected brain areas whose connectivity significantly tracked changes of one or more affective dimensions at all explored timescales. This conjunction analysis can be summarized as follows:

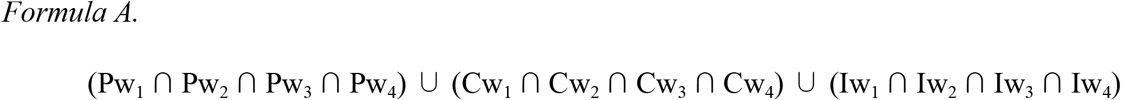

where Pw_i_, Cw_i_ and Iw_i_, represent regions significantly encoding, the polarity, complexity and intensity at the i-th timescale. This entire procedure was repeated for workflow A and B. To control for false positives, we pooled together p-values obtained from both workflows and all affective dimensions and applied the false discovery rate algorithm under dependency assumptions (Benjamini and Yekutieli, 2001; p-value < 0.05).

## Results

### Experiment 1: Single-Subject Affective Dimensions obtained from Behavioral Ratings

Affective states elicited by watching Forrest Gump are described by three orthogonal affective dimensions (Lettieri et al., 2019): (a) polarity defines whether the current experience is perceived as pleasant (negative scores) or unpleasant (positive scores); (b) complexity represents cognitively mediated states (positive scores) and instinctual responses (negative scores); (c) intensity denotes the emotional impact of the experience (monotonic increase in positive scores; for further details, see Supplementary Data).

Results of the estimate of the optimal window width show that a tradeoff between the across-participants agreement and window size is reached at 273 (9m:6s), 239 (7m:58s), 143 (4m:46s) and 89 timepoints (2m:58s) for polarity; also, the optimal window size is 205 (6m:50s), 143 (4m:46s), 109 (3m:38s) and 88 timepoints (2m:56s) for complexity and 331 (11m:2s), 247 (8m:14s), 163 (5m:26s) and 100 timepoints (3m:20s) for intensity.

### Experiment 1: Association between tISFC and Polarity of the Emotional Experience

The connectivity strength of three regions is significantly associated with changes in the polarity of the affective experience at all explored timescales (p_FDR_ < 0.05; Figure 2A; Supplementary Table 1). These three regions are the right dorsal prefrontal cortex (R-dPFC) pertaining to the control network, the right insula (R-INS), a node of the ventral attention network and the left precuneus (L-preCUN), a node of the DMN. Additionally, connectivity of three other DMN regions, the right temporo-parietal junction (R-TPJ), the right precuneus (R-preCUN) and the right dorsomedial prefrontal cortex (R-dmPFC), encodes polarity at three out of four timescales. Other brain areas, such as the right middle frontal gyrus (R-MFG), the right orbitofrontal cortex (R-OFC), the right postcentral gyrus (R-postCG), the left mid-cingulate cortex (L-mCING), the left middle occipital gyrus, the left insula and the left central sulcus, are significantly associated with changes in polarity at two or only one time window. Except for L-MOG, all other significant regions are negatively associated with polarity scores, meaning that the more unpleasant the experience is (i.e., positive scores), the weaker the connectivity of these nodes with the rest of the brain (Supplementary Figure 1A).

Overall, connectivity strength of DMN and control network nodes plays a major role in representing the pleasantness of the emotional experience during naturalistic stimulation (Figure 2A, Supplementary Table 1).

### Experiment 1: Association between tISFC and Intensity of the Emotional Experience

Findings demonstrate that connectivity strength of four regions is associated with the perceived intensity of the emotional experience (p_FDR_ < 0.05; Figure 2B; Supplementary Table 2). These four parcels are the right ventral prefrontal cortex (R-vPFC), a node of the limbic network, the R-TPJ - pertaining to the DMN -, the right superior frontal gyrus (R-SFG), belonging to the control network and the left amygdala (L-Amyg). Intensity is also related to connectivity of the right precentral gyrus (R-preCG), of the posterior portion of the right supramarginal gyrus (R-SMG) and of the right superior parietal lobule (R-SPL), at three out of four time windows. These brain regions pertain to the control, dorsal attention, and somatomotor networks. Other areas, such as the R-OFC, the right dorsal and ventral precentral sulcus (R-dpreCS and R-vpreCS, respectively), the R-INS, the right medial superior frontal gyrus (R-mSFG), the ascending ramus of the right cingulate sulcus (R-arCING), the left precentral gyrus (L-preCG), the left postcentral sulcus (L-postCS), the left fusiform gyrus (L-FusG) and the left hippocampus (L-Hipp) encode intensity at two or only one timescale. Except for the bilateral precentral gyrus, the L-FusG, the L-Amyg and L-Hipp, all other significant regions are positively associated with intensity values, meaning that the more intense the experience is, the stronger the connectivity of these nodes with the rest of the brain (Supplementary Figure 2A). Altogether, connectivity strength of control and dorsal attention networks plays a major role in tracking the intensity of the emotional experience (Figure 2B, Supplementary Table 2).

### Experiment 1: Association between tISFC and Complexity of Emotional Experience

No brain region is significantly associated with changes in the complexity dimension at all explored time windows (Supplementary Figure 3 and 5; Supplementary Table 3). Also, results show that connectivity strength of the right mid-cingulate cortex (R-mCING), a node of the somato-motor network, relates to complexity at three out of four timescales. Two parcels within the right inferior temporal gyrus (R-ITG), pertaining to the control and the dorsal attention networks, the right intraparietal sulcus (R-IPS), also included in the control network, and the left dorsal prefrontal cortex (L-dPFC), a node of the DMN, track changes in complexity at one timescale (Supplementary Figure 4). Concerning the interpretation of these relationships, R-ITG and R-IPS increase the strength of their connectivity with the rest of the brain during cognitively mediated states (e.g., mixed emotions; Lettieri et al., 2019), whereas R-mCING and L-dPFC are more strongly connected during movie scenes eliciting more instinctual responses (e.g., fear).

### Experiment 2: Validation of the relationship between tISFC and Affective Dimensions

To test the reliability and generalizability of results obtained from Forrest Gump (i.e., exploratory dataset), we collected and analyzed emotion dimension ratings of the live-action movie 101 Dalmatians (i.e., validation dataset; for further details, please see Supplementary Data). We measured the association between connectivity strength and affective dimensions after restricting the search for significance to regions encoding at least one dimension in the exploratory dataset at all explored timescales. This set of brain areas includes the R-TPJ, the R-SFG, the R-INS, the R-dPFC, the R-vPFC, the L-preCUN and the L-Amyg (Figure 3A). As no brain region encodes complexity at all explored timescales, none of the selected regions is obtained from the analysis of this dimension. Therefore, validation analyses are limited to polarity and intensity. Moreover, the selected regions pertain to the default, the control, the limbic, the ventral attention, and the subcortical networks.

**Figure 3.**
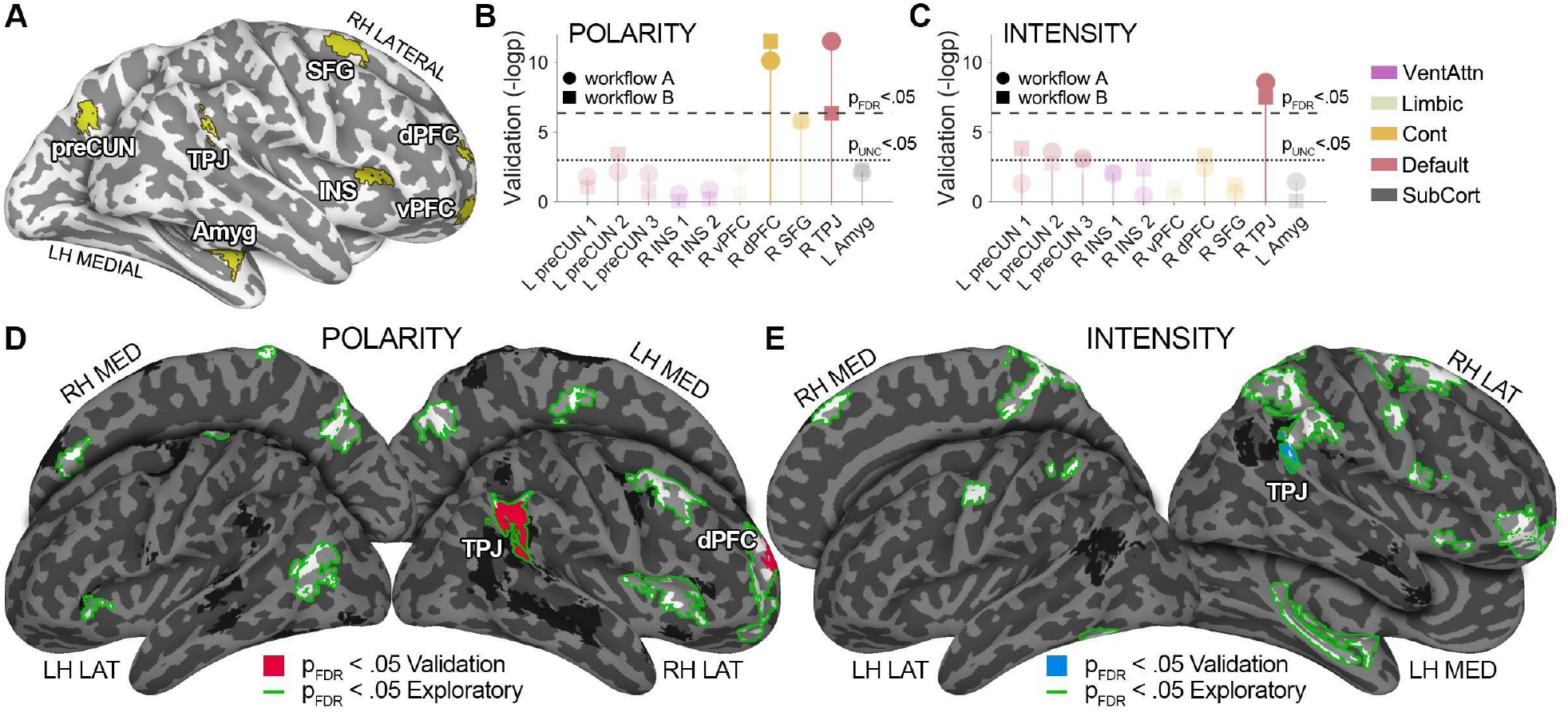
shows the results of the validation dataset. Panel **A** depicts the set of regions whose connectivity is associated with either polarity or intensity at all explored timescales in the exploratory dataset (i.e., conjunction analysis, see Formula A). The Manhattan plot in panel **B** and **C** summarizes the significance of the association between tISFC and affective dimensions in the validation dataset (i.e., -logp), across all regions obtained from experiment 1. Connectivity of the right temporo-parietal junction, a relevant node of the default mode network, is associated with changes in polarity and intensity, irrespectively of the fMRI preprocessing pipeline (p_FDR_<0.05 for workflows A and B). Also, connectivity of the right dorsal prefrontal cortex (control network) is associated with changes in polarity (p_FDR_<0.05 for workflows A and B). Panel **D** and **E** show whole-brain results. Green outlines delineate significant regions in the exploratory dataset; red (polarity; panel **D**) and blue fill (intensity; panel **E**) highlight regions associated with affect in the validation dataset after applying whole-brain FDR correction. Findings obtained from the whole-brain approach confirm ROI-based ones. LH = left hemisphere; RH = right hemisphere; Amyg = amygdala; preCUN = precuneus; TPJ = temporoparietal junction; INS = insula; dPFC = dorsal prefrontal cortex; vPFC = ventral prefrontal cortex; SFG = superior frontal gyrus; VentAttn = ventral attention network; Cont = control network; SubCort = subcortical network.

Results show that, in the validation dataset, both polarity and intensity ratings are associated with connectivity strength of R-TPJ, a DMN node (p_FDR_ < 0.05; Figure 3B-E). Importantly, this finding does not depend on the fMRI data preprocessing pipeline, and the relationship between affect and brain connectivity is interpreted as in the exploratory dataset: the more pleasant and intense the experience is, the stronger is the connectivity between this DMN area and the rest of the brain (Supplementary Figure 6 and 7).

In addition, connectivity strength of R-dPFC, a control network region, is significantly associated with changes in polarity in the validation dataset (p_FDR_ < 0.05 for both workflow A and B; Figure 3B and 3D). Similarly to R-TPJ, the interpretation of the relationship between R-dPFC connectivity and polarity is identical across the two experiments: the more pleasant the experience is, the more R-dPFC is connected to the rest of the brain (Supplementary Figure 6 and 7).

In summary, results of the validation dataset demonstrate that the relationship between polarity and connectivity of R-dPFC (control network), as well as the association between R-TPJ (DMN) and both polarity and intensity generalize to another movie, sample, different fMRI preprocessing workflows and setup for the collection of behavioral data.

### Experiment 1: Chronotopic Maps of Affective Dimensions

Here, we show the preferred timescale at which connectivity of significant regions relates to changes in polarity, complexity, and intensity. This analysis reveals the existence of chronotopic maps (Protopapa et al., 2019), where adjacent cortical regions preferentially map the stream of affect at different time intervals (Figure 4).

**Figure 4.**
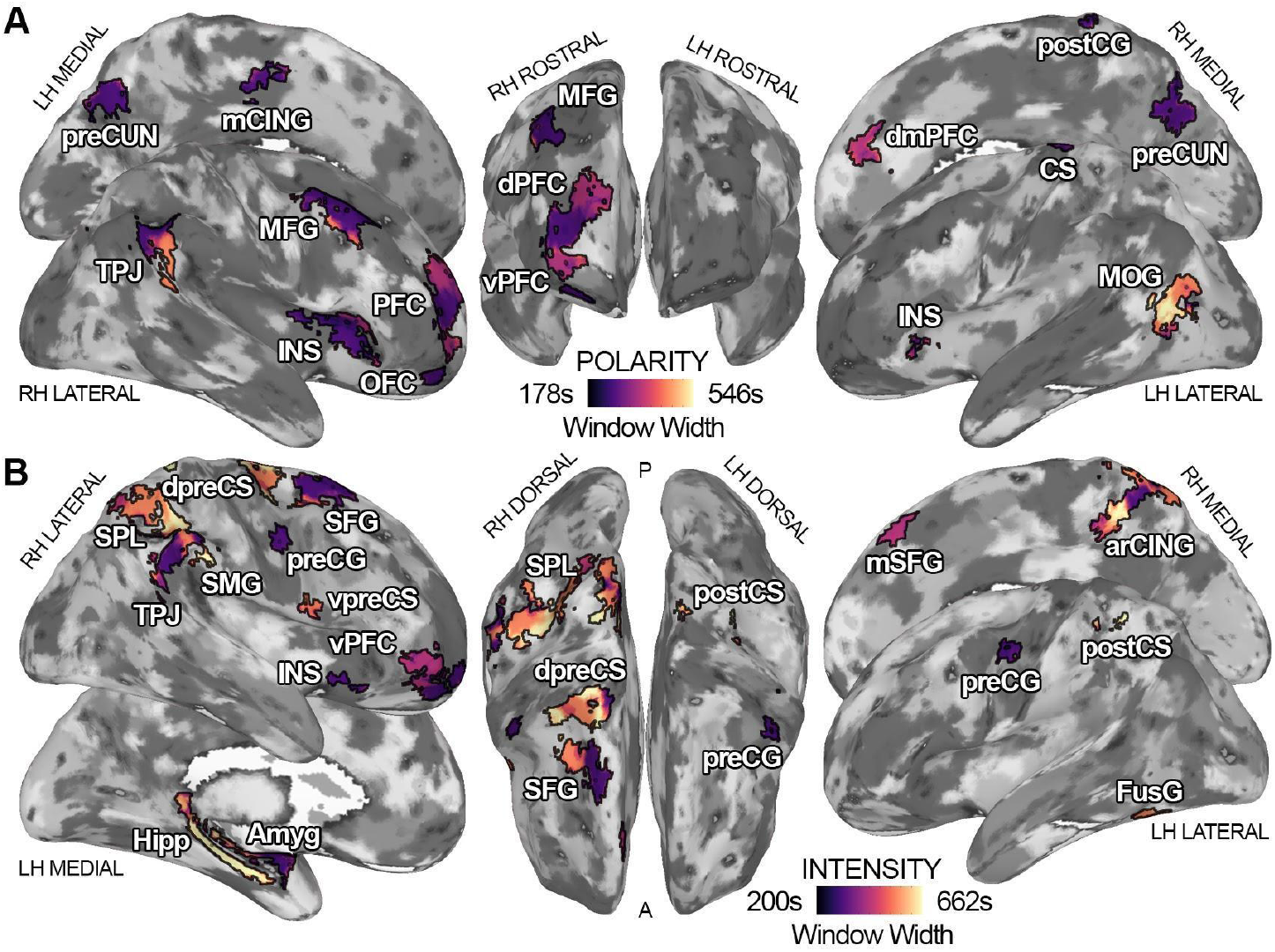
depicts the timescale at which connectivity dynamics are maximally associated with changes in polarity (panel **A**) and intensity (panel **B**) for each brain region. Dark purple indicates a preference for shorter timescales (∼178s for polarity and ∼200s for intensity), whereas bright yellow signals a preference for the mapping of affect in longer intervals (∼546s for polarity and ∼662s for intensity). LH = left hemisphere; RH = right hemisphere; Amyg = amygdala; preCUN = precuneus; TPJ = temporoparietal junction; INS = insula; dPFC = dorsal prefrontal cortex; vPFC = ventral prefrontal cortex; SFG = superior frontal gyrus; VentAttn = ventral attention network; Cont = control network; SubCort = subcortical network.

For the polarity dimension, chronotopic organizations are revealed in the right temporo-parietal and the right prefrontal cortex (Figure 4A). Within TPJ, connectivity of dorsocaudal territories is associated with polarity scores at shorter timescales (∼3 minutes), while ventrorostral portions preferentially track polarity in ∼8-minute segments. In the right prefrontal cortex, connectivity is mainly associated with scores of the polarity component at shorter timescales (∼3 minutes) in its central region, and at intermediate intervals (∼5 minutes) in its ventral and dorsal parts. In addition, connectivity of the R-dmPFC preferentially represents polarity in 5-minute segments, whereas L-MOG shows a marked preference for longer timescales (from ∼8 to ∼9 minutes). Connectivity of all other brain areas is maximally associated with this affective dimension at shorter intervals (∼3 minutes).

For the intensity component, we show timescale topographies within the right parietal lobe, the right frontopolar cortex, the right cingulate region and the R-SFG (Figure 4B). In the right parietal lobe, territories of the R-SPL preferentially map the intensity of the emotional experience in longer segments (from ∼8 to ∼11 minutes), whereas regions pertaining to the inferior parietal lobule in shorter time windows (from ∼3 to ∼5 minutes). The chronotopic arrangement in the right frontopolar region indicates that the connectivity of its lateral portion preferentially encodes intensity in ∼5-minute segments, whereas its medial part in ∼3-minute intervals. Also, within the right cingulate region, we observe an abrupt change in timescale preference, with connectivity of the inferior portion of the R-arCING preferentially mapping emotional intensity at the longest timescale (∼11 minutes) and its superior region at the shortest one (∼3 minutes). For the R-SFG, the medial part optimally represents intensity in ∼3-minute segments and the lateral portion in ∼8-minute ones. Interestingly, connectivity of L-Hipp and L-Amyg shows opposite behaviors: the hippocampus encodes emotional intensity at the longest timescale (∼11 minutes), whereas the amygdala at the shortest one (∼3 minutes). All other brain regions, besides the R-dpreCS, the L-postCS and the L-FusG are tuned to shorter time windows.

Lastly, no region significantly associated with the complexity dimension shows a chronotopic organization (Supplementary Figure 5).

## Discussion

In the current study, we combined real-time reports of the affective experience with recordings of fMRI activity during naturalistic stimulation to reveal how brain connectivity encodes the stream of affect. We collected approximately 45 hours of behavioral reports and 38 hours of brain imaging data over two experiments. In experiment 1, ratings of emotion categories associated with the watching of Forrest Gump were used to obtain timeseries of affective dimensions and to explain brain connectivity dynamics in independent individuals. In experiment 2, the 101 Dalmatians movie was used to test the reliability and generalizability of results obtained from the exploratory dataset: moment-by-moment scores of the polarity and intensity of the emotional experience were correlated with changes in the connectivity of brain regions identified in experiment 1. Also, fMRI data were preprocessed according to two alternative workflows. Therefore, our study protocol ensures that findings cannot be merely ascribed to (1) the selection of a specific movie, (2) participants’ characteristics, (3) the method used to collect behavioral reports, (4) the pipeline adopted to preprocess fMRI data.

Results show that connectivity strength of the right temporoparietal junction and of the right dorsal prefrontal cortex, respectively DMN and control network nodes, is positively associated with the polarity of the affective experience. Moreover, affective dimensions converge in the right temporo-parietal cortex, as this region’s connectivity tracks changes in emotional intensity as well. Together with the right prefrontal cortex, the right temporo-parietal junction represents the dynamics of affect in a chronotopic manner, meaning that adjacent portions of these areas preferentially map the stream of affect at different time intervals, ranging from few to several minutes.

In everyday life, affect varies as a function of time, and this complex relationship is hardly recreated in the laboratory setting. To elicit a variety of affective states, track their evolution in time and maintain a relatively high ecological validity, an optimal solution may be represented by movie watching. Movies produce strong affective responses over a few hours and mimic everyday life situations with an alternation of sudden and predictable events (Schaefer et al., 2010). Also, changes in affect induced by movies are based on rich contextual information and are related to the viewer’s expectations. Building upon this, we recorded brain activity and affective ratings during the watching of two movies, and demonstrated that changes in the subjective emotional experience explain brain connectivity dynamics in independent participants.

Over the last decade, researchers have emphasized the importance of moving from a region- to a network-based approach when studying the neural correlates of emotion (Lindquist and Barrett, 2012; Raz et al., 2016; Kragel et al., 2016; Pessoa, 2017). Changes in affect are associated with modifications in bodily responses (e.g., autonomic, musculoskeletal) and mental processes (e.g., memory, decision making), which are likely reflected in a distributed functional reorganization of brain modules (Pessoa, 2017). In this regard, the DMN is thought to play a crucial role in interoception (Kleckner et al., 2017) and in constructing and representing emotions (Satpute and Lindquist, 2019). Connectivity of this network is also altered in affective disorders, such as anxiety (Liao et al., 2010) or depression (Whitfield-Gabrieli and Ford, 2012). Our findings support the central role of this network in the processing of affect, as we demonstrate that its connectivity tracks changes in the subjective judgment of emotional polarity and intensity in a naturalistic environment. Furthermore, we show across two experiments that connectivity of a prefrontal control network node is also related to the polarity of affect. The control network is involved in empathy (Christov-Moore et al., 2020) and seems crucial in selecting sensations for conscious awareness during the construction of a mental state (Lindquist and Barrett, 2012). In our study, participants of the behavioral experiment were asked to constantly report their conscious emotional experience, indeed, a task that involves the processing and selection of sensations. It is interesting to note that, although fMRI participants were not explicitly asked to focus on their emotions, watching an evocative movie modulates connectivity of the control network, and such modulation is associated with conscious reports of affect.

When we look closely at nodes encoding affective dimensions, we notice that connectivity between the right temporoparietal junction and the rest of the brain represents both polarity and intensity. This cortical area is one of the core regions of the social brain (Saxe and Kanwisher, 2003; Lamm et al., 2007; Kohn et al., 2014; Etkin et al., 2015) and represents affective dimensions through orthogonal and spatially overlapping gradients (Lettieri et al., 2019). In our previous study, we hypothesized that this region could also act as a central node of a distributed network. Indeed, here we confirm our prediction by demonstrating that connectivity between temporo-parietal cortex and the rest of the brain tracks multiple descriptions of affect over time.

In experiment 1, we characterize the preferred timescale at which brain connectivity encodes the stream of affect. Previous studies on memory and narrative comprehension demonstrate that while sensory regions extract information in short segments, high-order areas preferentially encode longer events (Hasson et al., 2015; Simony et al., 2016; Baldassano et al., 2017). Here, we show that chronotopic maps (Protopapa et al., 2019) represent the temporal dynamics of the emotional experience at multiple timescales. Indeed, behavioral reports of affect are characterized by relatively slow dynamics, approximately between 3 and 11 minutes, as revealed by the data-driven estimate of the optimal timescale. This likely indicates that variations in affect during movie watching are associated with major changes in the plot. Accordingly, regions encoding affect are mainly high-order transmodal cortical areas, which are tuned to longer timescales (Baldassano et al., 2017). In addition, the right temporoparietal junction and the right prefrontal cortex show peculiar topographies when considering the preferred timescale. For polarity, the timescale increases moving from dorsocaudal to ventrorostral territories; instead, the right prefrontal cortex has a patchier organization, with its ventral and dorsal parts representing polarity at intermediate timescales and the central portion at shorter intervals. For intensity, the topography revealed in the right parietal lobe resembles the one reported for event structure in narratives (Baldassano et al., 2017), with connectivity of superior parietal regions encoding slower dynamics and inferior parietal areas faster ones.

It should be noted that we assessed the chronotopic organization of affect in the first experiment only, as this analysis requires exploring a wide range of temporal dynamics (i.e., from a few to tens of minutes). This was ensured by the movie we selected for experiment 1, as it lasts twice as long as the one employed in experiment 2. Further studies could test the reproducibility of chronotopic connectivity maps of affect using stimuli comparable in overall duration and with similar dynamics in emotional ratings.

In conclusion, here, we demonstrate that brain connectivity dynamics track changes in subjective reports of the affective experience. Specifically, the right temporo-parietal and the right dorsal prefrontal cortex, nodes of the DMN and control network, represent the polarity of the experience. Moreover, polarity and intensity converge into the right temporoparietal cortex, as connectivity of this region tracks the emotional impact as well. Within the right temporo-parietal junction and the right prefrontal cortex, the stream of affect is also represented at multiple timescales following chronotopic connectivity maps. Altogether, these findings reveal that the human brain represents changes in affect through connectivity of the default mode and control networks and that the timescale of emotional experiences is topographically mapped.

## Acknowledgments

We would like to thank all the people behind the studyforrest project, especially Michael Hanke. G.L., G.H. and L.C. are supported by Progetto di Attività Integrate - PAI Project - granted by IMT School for Advanced Studies Lucca. M.D. is supported by an ERC Consolidator Grant 2017 (LIGHTUP, project # 772953). L.C. would like to thank Fondazione IRIS for their support and funding.

## Author Contributions

G.L., G.H., P.P. and L.C., conceived the study. G.L., G.H. and L.C. designed the behavioral experiment, collected behavioral data, developed the code, performed behavioral and fMRI data analysis, interpreted the obtained results and drafted the manuscript. E.M.C. acquired behavioral data of the validation dataset. F.S., V.B., M.D. and A.L. acquired fMRI data of the validation dataset. G.L., G.H., L.C., F.S. and A.L. analyzed fMRI data of the validation dataset. G.L., G.H., E.M.C., F.S., V.B., M.D., A.L., E.R., P.P. and L.C. critically revised the manuscript. All the authors approved the final version of the manuscript.

