## Supplementary Data for "Default and Control networks connectivity dynamics track the stream of affect at multiple timescales"

**Supplementary Methods**


1. *Experiment 1: Behavioral Study - Participants.*
2. *Experiment 1: Behavioral Study - Single-Subject Affective Dimensions.*
3. *Experiment 1: fMRI Study - Participants.*
4. *Experiment 1: fMRI Study - Data Pre-processing.*
5. *Experiment 2: Behavioral Study - Stimuli and Experimental Paradigm.*
6. *Experiment 2: Behavioral Study - Single-Subject Affective Dimensions.*
7. *Experiment 2: fMRI Study - Participants.*
8. *Experiment 2: fMRI Study - Data Analysis.*
9. *Experiment 2: fMRI Study - Association between tISFC and Changes in Affective Dimensions.*


**Supplementary Results**


1. *Experiment 1: Single-Subject Affective Dimensions obtained from Behavioral Ratings.*
2. *Experiment 2: Validation of the relationship between tISFC and Affective Dimensions.*


**Supplementary Figures and Tables**

**Supplementary References**

**Supplementary Methods**

**1.** *Experiment 1: Behavioral Study - Participants.* All participants gave their written informed consent to take part in the study after risks and procedures had been explained. They retained the right to withdraw at any time and received a small monetary compensation for their participation. All subjects were clinically healthy and had no history of any neurological or psychiatric condition. They also had normal hearing, normal or corrected to normal vision and reported no history of drugs or alcohol abuse.

**2.** *Experiment 1: Behavioral Study - Single-Subject Affective Dimensions.* Studies on perception demonstrated that such dimensions are preferentially encoded in topographic maps (Harvey et al., 2013) and this may be the case also for the correlates of affective experiences (Lettieri et al., 2019). For this reason, by using principal component analysis (PCA) and procrustes rotation, we derived in each subject the timecourse of the affective dimensions explaining behavioral reports of the emotional experience. In brief, timeseries representing the emotional experience of each subject were downsampled to 2-second resolution (3,600 timepoints), matching the temporal characteristics of the fMRI acquisition. Data were then averaged across subjects and PCA was used to reveal group-level affective dimensions. PCA was also performed on single-subject data and the results were aligned to best approximate group-level dimensions using procrustes orthogonal linear transformation (reflection and rotation only). We considered the rotated scores of principal components as single-subject affective dimensions and used these data to inform subsequent fMRI analyses. To assess the agreement across subjects in affective dimensions, we also computed pairwise Spearman's correlation and its significance (see Lettieri et al., 2019 for further details).

**3.** *Experiment 1: fMRI Study - Participants.* Brain data comprise recordings of fourteen healthy German subjects (6F; mean age 29.4 years, range 20–40 years), instructed to simply inhibit any movement and enjoy the movie. Similarly to the behavioral part of the study, the fMRI acquisition lasted two hours and was split in eight runs (3T Philips Achieva scanner; 32 channels head coil; gradient recall echo-echo planar imaging; 2000ms repetition time, 30ms echo time, 90° flip angle, 3 mm isotropic voxel, 240 mm field of view). Together with the fMRI data, 3D T1w high-resolution anatomical images were also acquired.

**4.** *Experiment 1: fMRI Study - Data Pre-processing.* AFNI v.17.2.00 (Cox, 1996) and ANTs (Avants et al., 2009) were used to preprocess the MRI data. For each subject, structural images were first skull-stripped (*antsBrainExtraction.sh*) and transformed to match the MNI152 template using non-linear registration (*3dQwarp*). Functional data were corrected for intensity spikes (*3dDespike*), adjusted for slice timing acquisition (*3dTshift*) and corrected for head motion (*3dvolreg*) by computing the displacement between each volume and the most stable timepoint of each run (i.e., the one showing smallest framewise displacement values; Power et al., 2012). Afterwards, the *align_epi_anat.py* and *3dQwarp* software were used to estimate the co-registration between functional and structural data, as well as to correct for phase distortion by allowing non-linear deformations in the y-direction (i.e., phase acquisition direction) only. Linear (i.e., motion correction, co-registration) and non-linear (i.e., phase distortion and MNI152 registration) transformations were then concatenated and applied (*3dNwarpApply*) to the functional data, so to obtain standard-space single-subject timeseries with a single interpolation step (i.e., sinc interpolation method), maintaining also the original spatial resolution (3mm isotropic voxel). In addition, data were iteratively smoothed until 6mm full-width at half maximum level was reached (*3dBlurToFWHM*) and rescaled (*3dcalc*) so that changes in activity were expressed as percentage with respect to the average voxel intensity in time (*3dTstat*). Lastly, we used *3dDeconvolve* to regress out from brain activity signal drifts (*polort* in *3dDeconvolve*) and physiological confounds following the recommendations by Ciric et al., 2017. Therefore, we used a model with 36 regressors of no interest to reduce the possibility that results of intersubject functional correlation could be explained by factors other than neurovascular coupling. Specifically, we created (*Atropos*;

*3dmask_tool*) binary masks of the white matter (WM) and cerebrospinal fluid (CSF) and extracted the timeseries of the average WM and CSF activity. Global signal (GS), which is the average activity across all voxels, was also calculated. For each of the three obtained timeseries (i.e., GS, WM, CSF), we computed their temporal derivatives, the quadratic terms and squares of derivatives, resulting in 12 regressors. The remaining 24 comprised the six head motion parameters estimated with *3dvolreg*, their temporal derivatives, their quadratic terms and the squares of derivatives. All regressors of no interest were then de-trended using the same degree of polynomial (*polort*) employed in *3dDeconvolve* to remove slow drifts from voxels activity. The model was then fitted in brain activity using a mass-univariate general linear model. The residuals of this fitting were considered the single-subject brain activity associated with the watching of Forrest Gump and used in subsequent analyses.

**5.** *Experiment 2: Behavioral Study - Stimuli and Experimental Paradigm.* A shortened version of 101 Dalmatians was selected as a validation dataset. Similarly to what Hanke and colleagues (2016) did in Forrest Gump (i.e., exploratory dataset), we discarded scenes irrelevant to the central plot, so as to maintain the total running time of the movie below 1 hour (i.e., 54 minutes). Video editing was carried out using the iMovie software (10.1.10) on an Apple Macbook Air. The movie was also split into six runs of similar duration and a six-second fade-in and fade-out period was added at the beginning and the end of each run.

In the behavioral experiment subjects sat comfortably in a silent room facing a 24" Dell™ screen, wore headphones (Marshall™ Major III; 20–20,000 Hz; Maximum SPL 97 dB), and were asked to report their affective experience while watching 101 Dalmatians. As for the exploratory dataset, all participants have not watched the movie in the period of at least one year before the experiment. Stimulus presentation and recording of the responses were implemented in Matlab (R2019b; MathWorks Inc., Natick, MA, USA) and Psychtoolbox v3.0.16 (Kleiner et al., 2007).

**6.** *Experiment 2: Behavioral Study - Single-Subject Affective Dimensions.* For the exploratory dataset, we collected the perceived intensity of six emotions and then used PCA to obtain timeseries of polarity and intensity. In the validation dataset, instead, we modified the collection of behavioral reports, focusing directly on affective dimensions. In fact, participants were asked to report moment-by-moment (i.e., 10Hz sampling rate; 16,140 timepoints) the pleasantness or unpleasantness and intensity of the experience on a continuous scale ranging from -100 (extremely negative) to +100 (extremely positive). For each subject, raw ratings were interpreted as a measure of polarity, whereas its absolute value (i.e., the emotional impact regardless of the valence of the experience) was considered a measure of intensity. Lastly, to assess the agreement across subjects of polarity and intensity, we computed pairwise Spearman's correlation and its significance.

**7.** *Experiment 2: fMRI Study - Participants.* Brain activity elicited by the watching of 101 Dalmatians was obtained from ten healthy Italian subjects (8F; 35±13 years), instructed to simply inhibit any movement and enjoy the movie. The fMRI acquisition lasted 1 hour and was split in six runs (3T Philips Ingenia scanner, Neuroimaging centre of NIT - Molinette Hospital, Turin; 32 channels head coil; gradient recall echo-echo planar imaging; 2000ms repetition time, 30ms echo time, 75° flip angle, 3 mm isotropic voxel, 240 mm field of view). Audio and visual stimulation were delivered through MR-compatible LCD goggles and headphones (VisualStim Resonance Technology, video resolution 800x600 at 60 Hz, visual field 30° × 22°, 5″, audio 30 dB noise-attenuation, 40 Hz to 40 kHz frequency response). Together with the fMRI data, 3D T1w high-resolution anatomical images were also acquired (magnetization-prepared rapid gradient echo; 7ms repetition time, 3.2ms echo time, 9° flip angle, 1mm isotropic voxel, 224 mm field of view). The fMRI study was approved by the Ethical Committee of the University of Turin (Protocol No. 195874/2019) and participants gave their written consent for the participation.

**8.** *Experiment 2: fMRI Study - Data Analysis.* Considering the high variability in results related to analytical choices (Botvinik-Nezer et al., 2020), here we tested two alternative pre-processing pipelines (i.e., workflow A and B) for the analysis of the validation dataset.

Specifically, for workflow A, we removed scanner-related noise by applying a spike removal procedure (*3dDespike*). Afterward, all volumes were temporally aligned (*3dTshift*) and corrected for head motion (*3dvolreg*) by computing the displacement between each volume and the most stable timepoint of the first run (i.e., the one showing smallest framewise displacement values; Power et al., 2012). We applied a spatial smoothing up to 6mm (Gaussian kernel; *3dBlurToFWHM*) and data were normalized (i.e., mean centering and percentage scaling). Moreover, we removed the trend in data by applying the Savitzky-Golay filtering (MATLAB function *sgolayfilt*, polynomial order: 3, frame length: 200 timepoints; Çukur et al., 2013). Runs were then concatenated, and multiple regression analysis was performed (*3dDeconvolve*) to remove artifacts in signal related to head motion (6 motion parameters and spike regression for timepoint characterized by framewise displacement above 0.3). The residuals of this fitting were considered the single-subject brain activity associated with the watching of 101 Dalmatians, they were nonlinearly (*3dQWarp*) registered to the MNI standard space (Fonov et al., 2009) and used in subsequent analyses.

As a further control of the robustness of our findings, in workflow B, we followed the exact same pre-processing pipeline used for the exploratory dataset (i.e., experiment 1).

**9.** *Experiment 2: fMRI Study - Association between tISFC and Changes in Affective Dimensions.* To evaluate the reliability and generalizability of the association between tISFC and the timecourse of affective dimensions reported in experiment 1, we tested whether brain connectivity dynamics explain changes in polarity and intensity in 101 Dalmatians. To this aim, we first obtained timeseries of affective dimensions from each subject and estimated the optimal window width based on the intersubject correlation in polarity and intensity ratings (from 20 to 112 timepoints, 33% of overlap). Similarly to experiment 1, we considered the point at which the intersubject correlation curve starts to flatten (i.e., knee point) as the optimal window size. Afterwards, we estimated the tISFC as the Pearson's correlation between each ROI of one subject and all the other ROIs of the other subjects, across all the windows. We then applied Fisher z-transformation and averaged correlation values across pairings of subjects to produce a group-level timeseries of brain connectivity related to the watching of 101 Dalmatians. Also, by summing the Z-transformed correlation values at each timepoint, we obtained a timeseries of connectivity strength for each ROI. As in experiment 1, group-level polarity and intensity timeseries were downsampled to the optimal window width using a moving-average procedure and they were correlated to connectivity strength of each ROI over time (Spearman's $\rho$ coefficient). To assess the significance of the association, we used a non-parametric permutation test based on timepoint shuffling of affective dimensions (200,000 iterations; minimum two-tailed p-value: 1.0e-5). Importantly, we restricted analyses to ROIs significantly tracking changes in polarity and intensity in experiment 1, considering all explored timescales (see Figure 3A). This entire procedure was repeated for each analysis workflow (see previous section). Lastly, to control for false positives in experiment 2, we applied FDR correction ($p < 0.05$; Benjamini and Yekutieli, 2001) after pooling together p-values obtained from each timescale, affective dimension, and workflow.

**Supplementary Results**

**1.** *Experiment 1: Single-Subject Affective Dimensions obtained from Behavioral Ratings.* For polarity scores, median Spearman's correlation across subjects is ρ=0.580 (95% CI: 0.560 - 0.615) with an interquartile range of ρ=0.124. For *complexity* scores, median Spearman's correlation across subjects is ρ=0.437 (95% CI: 0.408 - 0.462) with an interquartile range of ρ=0.124. Lastly, for intensity ratings, median Spearman's correlation across subjects is ρ=0.410 (95% CI: 0.391 - 0.423) with an interquartile range of ρ=0.089.

**2.** *Experiment 2: Validation of the relationship between tISFC and Affective Dimensions.* We first downsampled polarity and intensity behavioral reports to match the fMRI resolution (1,614 timepoints) and correlated the obtained timeseries across subjects. For polarity, median Spearman's correlation across subjects is ρ=0.487 (95% CI: 0.452 - 0.530) with an interquartile range of ρ=0.293. Also, the 97.1% (204 out of 210) of correlations across all possible pairings of subjects reach statistical significance after correction for multiple comparisons ($p_{Bonf} < 0.05$). For intensity, instead, median correlation is ρ=0.229 (95% CI: 0.203 - 0.245) with an interquartile range of ρ=0.184 and the 88.6% (186 out of 210) of correlation values are significant ($p_{Bonf} < 0.05$). As in experiment 1, we downsampled affective dimension timeseries based on the estimate of the optimal window size (polarity: 67 timepoints; intensity: 37 timepoints). Group averaged polarity and intensity ratings were then used to explain the time-varying brain connectivity in independent subjects watching the same movie. We restricted these analyses to regions significantly encoding affective dynamics in the exploratory dataset (i.e., experiment 1) and applied correction for multiple comparisons based on the number of regions of interest.

### Supplementary Figures and Tables

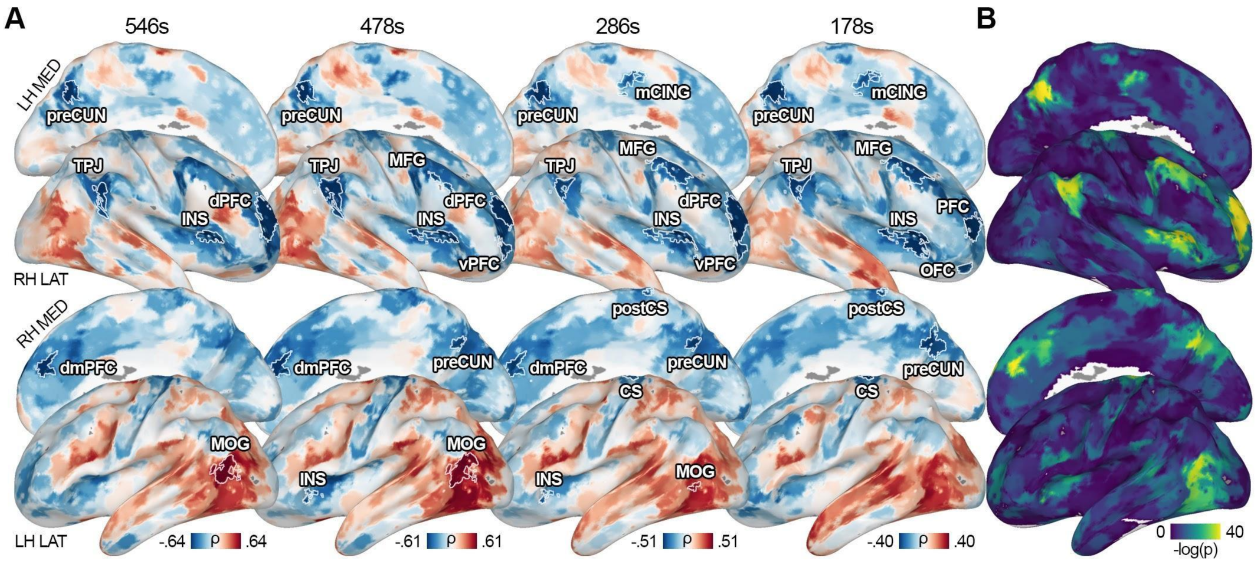


**Supplementary Figure 1.** Panel **A** shows results for the correlation (i.e., Spearman's rho) between polarity and brain connectivity dynamics at each timescale. Panel **B** represents the association between tISFC strength of each brain region and polarity across timescales. Colors reflect the sum of -log(p-value) across timescales, ranging from no involvement at any timescale (i.e., dark blue) to consistent recruitment (i.e., yellow). LH = left hemisphere; RH = right hemisphere; LAT = lateral; MED = medial; preCUN = precuneus; TPJ = temporoparietal junction; INS = insula; dPFC = dorsal prefrontal cortex; vPFC = ventral prefrontal cortex; mCING = mid cingulate cortex; MFG = middle frontal gyrus; dmPFC = dorsomedial prefrontal cortex; MOG = middle occipital gyrus; PFC = prefrontal cortex; CS = central sulcus; postCS = postcentral sulcus; OFC = orbitofrontal cortex.

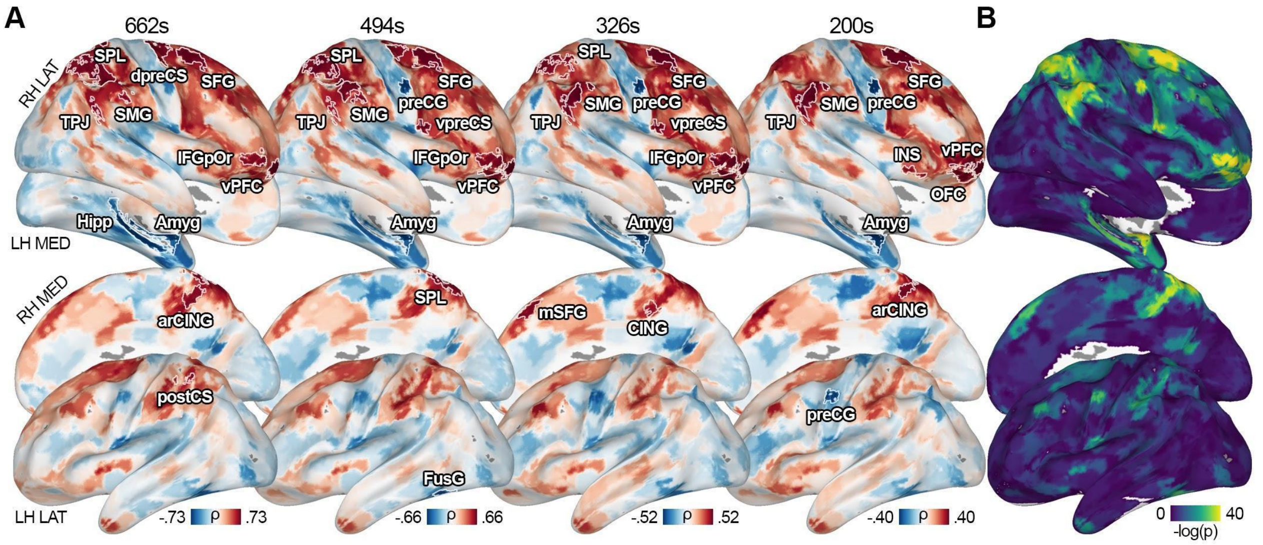


**Supplementary Figure 2.** Panel **A** shows results for the correlation (i.e., Spearman's rho) between intensity and brain connectivity dynamics at each timescale. Panel **B** represents the association between tISFC strength of each brain region and intensity across timescales. Colors reflect the sum of -log(p-value) across timescales, ranging from no involvement at any timescale (i.e., dark blue) to consistent recruitment (i.e., yellow). LH = left hemisphere; RH = right hemisphere; LAT = lateral; MED = medial; arCING = ascending ramus of the right cingulate sulcus; Amyg = amygdala; Hipp = hippocampus; TPJ = temporoparietal junction; INS = insula; vPFC = ventral prefrontal cortex; CING = cingulate cortex; FusG = fusiform gyrus; preCG = precentral gyrus; postCS = postcentral sulcus; dpreCS = dorsal precentral sulcus; vpreCS = ventral precentral sulcus; SFG = superior frontal gyrus; IFGpOr = inferior frontal gyrus pars orbitalis; mSFG = medial superior frontal gyrus; SPL = superior parietal lobule; SMG = supramarginal gyrus; OFC = orbitofrontal cortex.

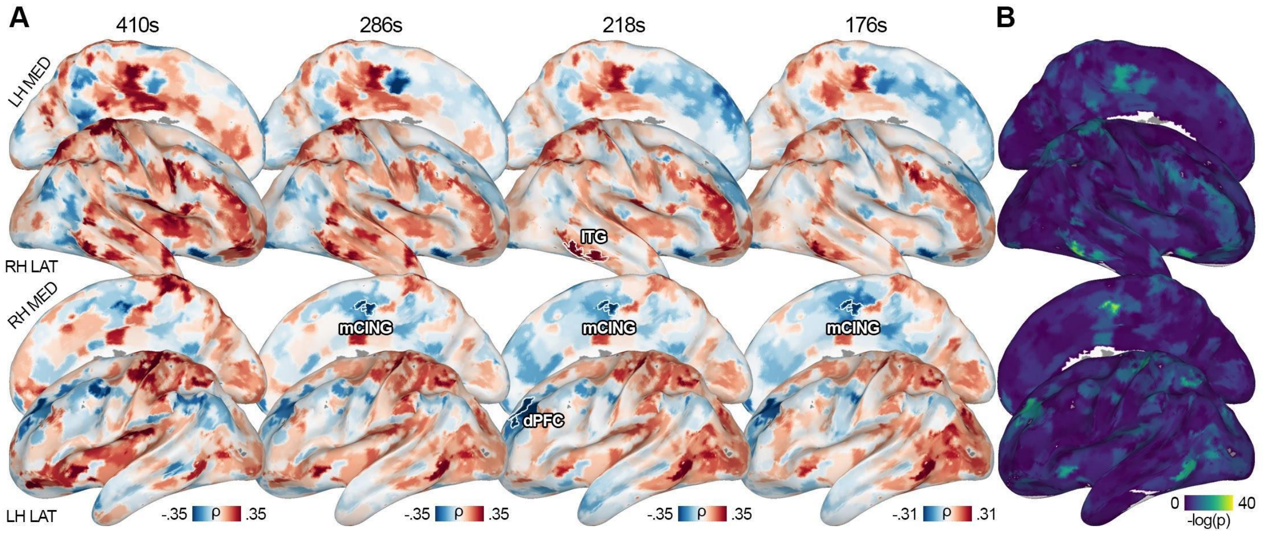


**Supplementary Figure 3.** Panel **A** shows results for the correlation (i.e., Spearman's rho) between complexity and brain connectivity dynamics at each timescale. Panel **B** represents the association between tISFC strength of each brain region and complexity across timescales. Colors reflect the sum of -log(p-value) across timescales, ranging from no involvement at any timescale (i.e., dark blue) to consistent recruitment (i.e., yellow). LH = left hemisphere; RH = right hemisphere; LAT = lateral; MED = medial; mCING = medial cingulate cortex; dPFC = dorsal prefrontal cortex; ITG = inferior temporal gyrus.

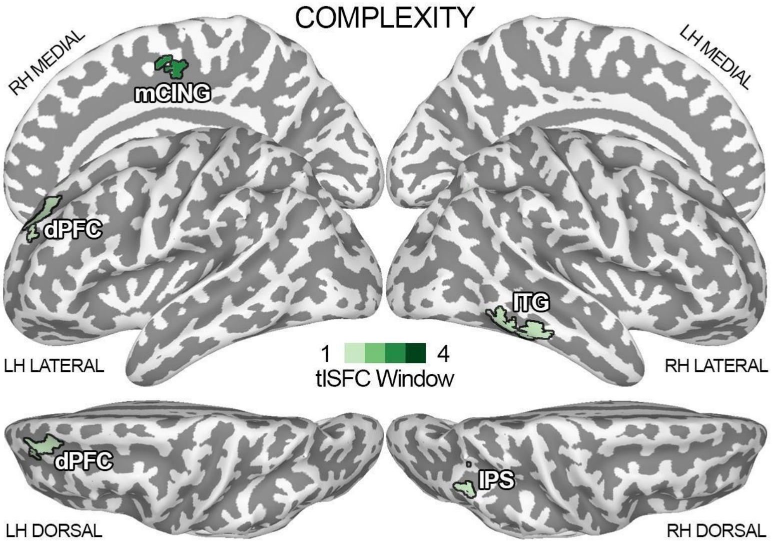


**Supplementary Figure 4.** This figure shows results for the connectivity strength of brain regions associated with changes in the complexity of the affective experience at all explored timescales. Grading of the color green indicates the number of window widths in which the connectivity strength of a region was associated with complexity. LH = left hemisphere; RH = right hemisphere; mCING = medial cingulate cortex; dPFC = dorsal prefrontal cortex; ITG = inferior temporal gyrus; IPS = intraparietal sulcus; tISFC = time-varying intersubject functional correlation.

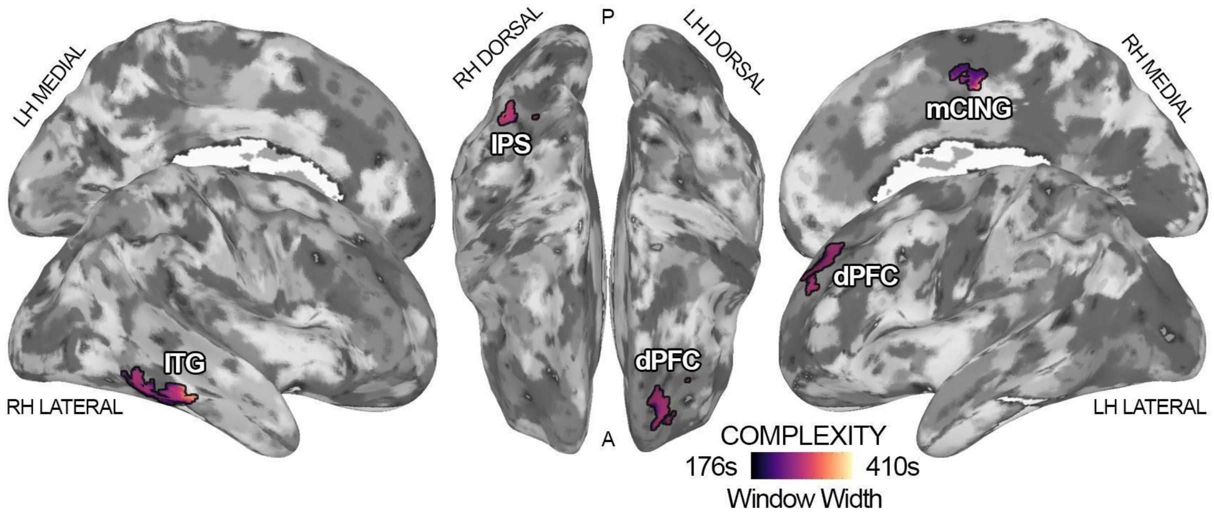


**Supplementary Figure 5.** This figure depicts the timescale at which connectivity dynamics are maximally associated with changes in complexity for each brain region. Dark purple indicates a preference for shorter timescales (~176s), whereas bright yellow signals a preference for the mapping of affect in longer intervals (~410s). LH = left hemisphere; RH = right hemisphere; mCING = medial cingulate cortex; dPFC = dorsal prefrontal cortex; ITG = inferior temporal gyrus; IPS = intraparietal sulcus.

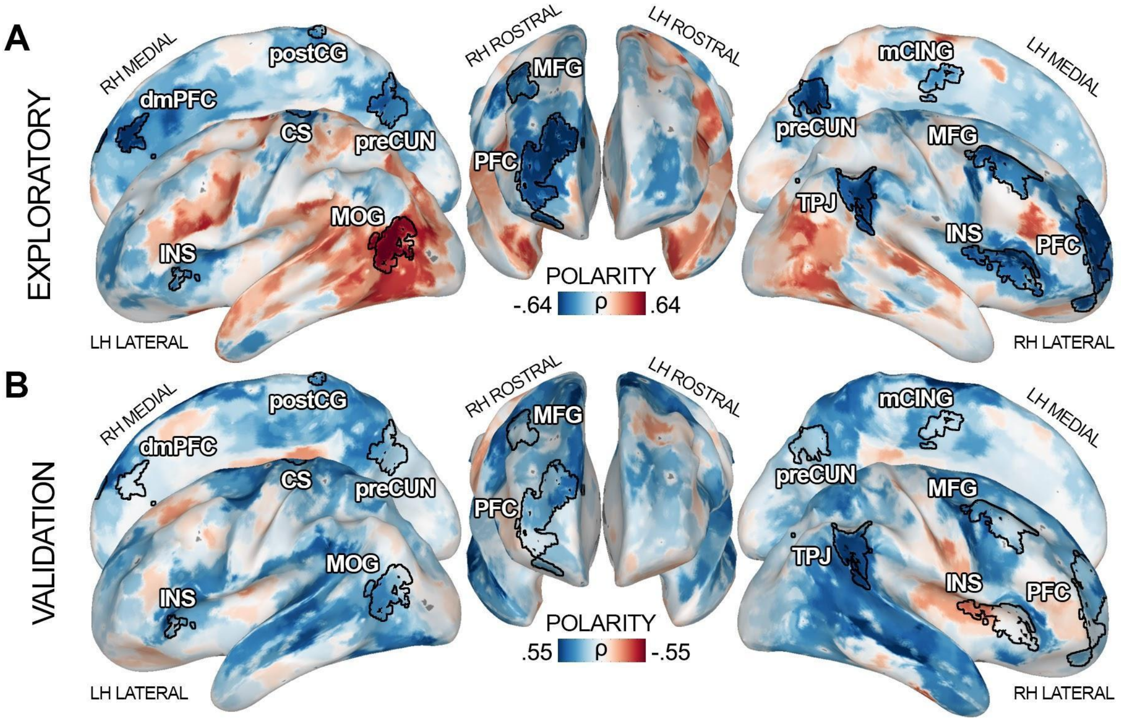


**Supplementary Figure 6.** This figure allows for a direct comparison of the exploratory (panel **A**) and the validation (panel **B**) datasets in terms of Spearman's correlation between brain connectivity and the timecourse of polarity. Please note that pleasant states were denoted by negative polarity scores in the exploratory dataset and positive in the validation one. Thus we inverted the colormap of the validation dataset to ease the direct comparison between the two experiments. LH = left hemisphere; RH = right hemisphere; preCUN = precuneus; TPJ = temporoparietal junction; INS = insula; mCING = mid cingulate cortex; MFG = middle frontal gyrus; dmPFC = dorsomedial prefrontal cortex; MOG = middle occipital gyrus; PFC = prefrontal cortex; CS = central sulcus; postCG = postcentral gyrus.

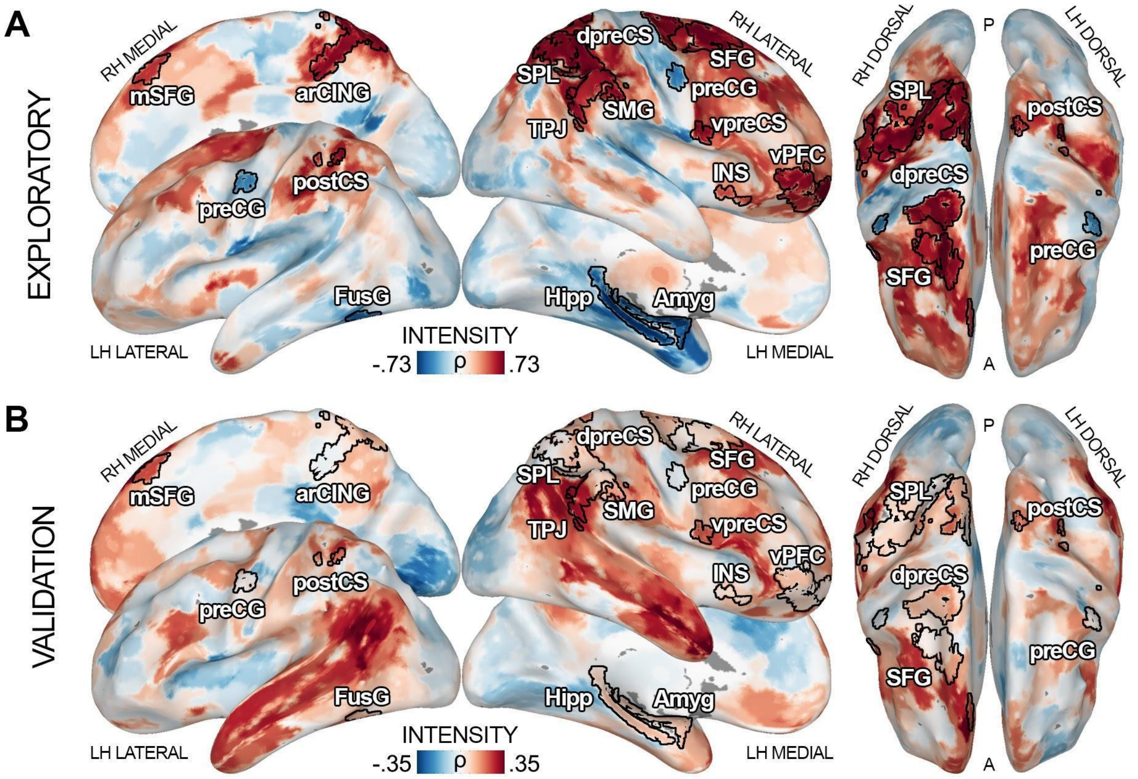


**Supplementary Figure 7.** This figure allows for a direct comparison of the exploratory (panel **A**) and the validation (panel **B**) datasets in terms of Spearman's correlation between brain connectivity and the timecourse of intensity. LH = left hemisphere; RH = right hemisphere; arCING = ascending ramus of the right cingulate sulcus; Amyg = amygdala; Hipp = hippocampus; TPJ = temporoparietal junction; INS = insula; vPFC = ventral prefrontal cortex; FusG = fusiform gyrus; preCG = precentral gyrus; postCS = postcentral sulcus; dpreCS = dorsal precentral sulcus; vpreCS = ventral precentral sulcus; SFG = superior frontal gyrus; mSFG = medial superior frontal gyrus; SPL = superior parietal lobule; SMG = supramarginal gyrus.

### Supplementary Table 1

| | POLARITY - EXPLORATORY DATASET | | TIMESCALES | | | | | |
|---|---|---|---|---|---|---|---|---|
| ROI | Region Name | Network | 273 | 239 | 143 | 89 | LPI Coord (x,y,z) | sum(-logp) |
| 876 | 7Networks_RH_Cont_PFCl_11 | Cont | • | • | • | • | 26,61,15 | 45.36 |
| 489 | 7Networks_LH_Default_pCunPCC_24 | Default | • | • | • | • | -2,-70,36 | 46.05 |
| 493 | 7Networks_LH_Default_pCunPCC_28 | Default | • | • | • | • | -5,-62,39 | 45.36 |
| 495 | 7Networks_LH_Default_pCunPCC_30 | Default | • | • | • | • | -3,-68,45 | 44.67 |
| 777 | 7Networks_RH_SalVentAttn_FrOperIns_12 | SalVentAttn | • | • | • | • | 34,20,10 | 44.26 |
| 778 | 7Networks_RH_SalVentAttn_FrOperIns_13 | SalVentAttn | • | • | • | • | 41,24,6 | 42.72 |
| 874 | 7Networks_RH_Cont_PFCl_9 | Cont | • | | • | • | 31,55,10 | 42.87 |
| 882 | 7Networks_RH_Cont_PFCl_17 | Cont | • | • | • | | 25,55,26 | 42.65 |
| 925 | 7Networks_RH_Default_Par_13 | Default | | • | • | • | 57,-53,33 | 38.52 |
| 966 | 7Networks_RH_Default_PFCdPFCm_10 | Default | • | • | • | | 16,64,21 | 42.01 |
| 967 | 7Networks_RH_Default_PFCdPFCm_11 | Default | • | • | • | | 8,45,22 | 40.52 |
| 995 | 7Networks_RH_Default_pCunPCC_15 | Default | | • | • | • | 4,-65,36 | 37.91 |
| 779 | 7Networks_RH_SalVentAttn_FrOperIns_14 | SalVentAttn | | • | • | • | 34,5,11 | 40.17 |
| 362 | 7Networks_LH_Cont_pCun_2 | Cont | | | • | • | -10,-73,39 | 35.78 |
| 363 | 7Networks_LH_Cont_pCun_3 | Cont | | • | | • | -4,-76,45 | 37.28 |
| 869 | 7Networks_RH_Cont_PFCl_4 | Cont | | • | • | | 32,59,-2 | 34.92 |
| 892 | 7Networks_RH_Cont_PFCl_27 | Cont | | | • | • | 41,23,47 | 32.34 |
| 895 | 7Networks_RH_Cont_PFCl_30 | Cont | | | • | • | 38,10,58 | 37.68 |
| 917 | 7Networks_RH_Default_Par_5 | Default | • | • | | | 56,-47,22 | 30.43 |
| 922 | 7Networks_RH_Default_Par_10 | Default | • | • | | | 53,-46,33 | 37.51 |
| 182 | 7Networks_LH_DorsAttn_Post_10 | DorsAttn | • | • | | | -49,-62,10 | 36.62 |
| 183 | 7Networks_LH_DorsAttn_Post_11 | DorsAttn | • | • | | | -48,-69,16 | 37.99 |
| 249 | 7Networks_LH_SalVentAttn_FrOperIns_5 | SalVentAttn | | • | • | | -35,30,-1 | 34.91 |
| 128 | 7Networks_LH_SomMot_47 | SomMot | | | • | • | -8,-9,48 | 32.57 |
| 171 | 7Networks_LH_SomMot_90 | SomMot | | | • | • | -12,-26,74 | 34.93 |
| 674 | 7Networks_RH_SomMot_93 | SomMot | | | • | • | 5,-41,70 | 33.69 |
| 847 | 7Networks_RH_Cont_Par_5 | Cont | | | | • | 55,-51,46 | 29.91 |
| 867 | 7Networks_RH_Cont_PFCl_2 | Cont | | | | • | 28,52,-14 | 32.66 |
| 890 | 7Networks_RH_Cont_PFCl_25 | Cont | | • | | | 47,16,39 | 28.40 |
| 893 | 7Networks_RH_Cont_PFCl_28 | Cont | | | | • | 33,30,47 | 35.06 |
| 900 | 7Networks_RH_Cont_pCun_1 | Cont | | | | • | 14,-68,35 | 30.67 |
| 902 | 7Networks_RH_Cont_pCun_5 | Cont | | | | • | 5,-74,46 | 34.86 |
| 953 | 7Networks_RH_Default_PFCv_7 | Default | | | • | | 42,26,-5 | 33.49 |
| 179 | 7Networks_LH_DorsAttn_Post_7 | DorsAttn | | | • | | -48,-65,2 | 32.65 |
| 180 | 7Networks_LH_DorsAttn_Post_8 | DorsAttn | | • | | | -54,-68,6 | 33.51 |
| 185 | 7Networks_LH_DorsAttn_Post_13 | DorsAttn | | • | | | -42,-71,23 | 31.73 |
| 773 | 7Networks_RH_SalVentAttn_FrOperIns_8 | SalVentAttn | | | | • | 33,24,2 | 33.06 |

### Supplementary Table 2

| | INTENSITY - EXPLORATORY DATASET | | TIMESCALES | | | | | |
|---|---|---|---|---|---|---|---|---|
| ROI | Region Name | Network | 331 | 247 | 163 | 100 | LPI Coord (x,y,z) | sum(-logp) |
| 899 | 7Networks_RH_Cont_PFCl_34 | Cont | • | • | • | • | 24,9,63 | 45.36 |
| 921 | 7Networks_RH_Default_Par_9 | Default | • | • | • | • | 57,-53,24 | 41.45 |
| 825 | 7Networks_RH_Limbic_OFC_14 | Limbic | • | • | • | • | 21,62,-10 | 44.95 |
| 1002 | 7Networks_LH_SubCort_Amyg | SubCort | • | • | • | • | -24,-5,-17 | 43.65 |
| 843 | 7Networks_RH_Cont_Par_1 | Cont | | • | • | • | 61,-45,32 | 38.62 |
| 844 | 7Networks_RH_Cont_Par_2 | Cont | | • | • | • | 59,-46,41 | 39.97 |
| 848 | 7Networks_RH_Cont_Par_6 | Cont | | • | • | • | 60,-32,50 | 42.87 |
| 868 | 7Networks_RH_Cont_PFCl_3 | Cont | | • | • | • | 40,52,-12 | 39.00 |
| 870 | 7Networks_RH_Cont_PFCl_5 | Cont | • | • | • | | 44,49,-4 | 41.89 |
| 715 | 7Networks_RH_DorsAttn_Post_31 | DorsAttn | • | • | • | | 36,-45,52 | 39.12 |
| 716 | 7Networks_RH_DorsAttn_Post_32 | DorsAttn | • | • | • | | 32,-59,58 | 38.22 |
| 723 | 7Networks_RH_DorsAttn_Post_39 | DorsAttn | • | • | • | | 24,-65,57 | 39.22 |
| 726 | 7Networks_RH_DorsAttn_Post_42 | DorsAttn | • | • | • | | 37,-48,62 | 40.88 |
| 628 | 7Networks_RH_SomMot_47 | SomMot | | • | • | • | 55,-4,43 | 39.45 |
| 857 | 7Networks_RH_Cont_Par_15 | Cont | • | • | | | 43,-49,47 | 37.43 |
| 858 | 7Networks_RH_Cont_Par_16 | Cont | • | • | | | 44,-39,44 | 39.76 |
| 922 | 7Networks_RH_Default_Par_10 | Default | | | • | • | 53,-46,33 | 36.82 |
| 704 | 7Networks_RH_DorsAttn_Post_20 | DorsAttn | • | • | | | 49,-36,55 | 35.86 |
| 718 | 7Networks_RH_DorsAttn_Post_34 | DorsAttn | • | • | | | 30,-51,52 | 32.84 |
| 734 | 7Networks_RH_DorsAttn_Post_50 | DorsAttn | • | • | | | 16,-56,67 | 37.70 |
| 735 | 7Networks_RH_DorsAttn_Post_51 | DorsAttn | • | • | | | 9,-51,70 | 40.23 |
| 760 | 7Networks_RH_SalVentAttn_TempOccPar_15 | SalVentAttn | • | • | | | 62,-29,39 | 35.20 |
| 788 | 7Networks_RH_SalVentAttn_FrOperIns_23 | SalVentAttn | | • | • | | 56,9,13 | 38.91 |
| 807 | 7Networks_RH_SalVentAttn_Med_16 | SalVentAttn | • | | | • | 6,-47,57 | 37.53 |
| 164 | 7Networks_LH_SomMot_83 | SomMot | • | • | | | -14,-49,73 | 34.32 |
| 662 | 7Networks_RH_SomMot_81 | SomMot | • | • | | | 28,-11,65 | 34.99 |
| 845 | 7Networks_RH_Cont_Par_3 | Cont | | • | | | 59,-38,39 | 35.10 |
| 849 | 7Networks_RH_Cont_Par_7 | Cont | | • | | | 55,-41,48 | 36.15 |
| 852 | 7Networks_RH_Cont_Par_10 | Cont | | • | | | 49,-44,49 | 27.30 |
| 853 | 7Networks_RH_Cont_Par_11 | Cont | | • | | | 48,-50,55 | 33.06 |
| 854 | 7Networks_RH_Cont_Par_12 | Cont | | • | | | 49,-39,47 | 33.87 |
| 867 | 7Networks_RH_Cont_PFCl_2 | Cont | | | | • | 28,52,-14 | 34.29 |
| 898 | 7Networks_RH_Cont_PFCl_33 | Cont | | | | • | 20,18,62 | 37.32 |
| 911 | 7Networks_RH_Cont_PFCmp_4 | Cont | | | • | | 4,38,47 | 29.65 |
| 948 | 7Networks_RH_Default_PFCv_2 | Default | | | | • | 31,20,-13 | 27.20 |
| 175 | 7Networks_LH_DorsAttn_Post_3 | DorsAttn | | • | | | -43,-54,-17 | 29.12 |
| 211 | 7Networks_LH_DorsAttn_Post_39 | DorsAttn | • | | | | -38,-42,57 | 27.35 |
| 709 | 7Networks_RH_DorsAttn_Post_25 | DorsAttn | • | | | | 37,-42,44 | 29.44 |
| 730 | 7Networks_RH_DorsAttn_Post_46 | DorsAttn | | • | | | 11,-64,63 | 37.56 |

| | | | | | | | | |
|---|---|---|---|---|---|---|---|---|
| 731 | 7Networks_RH_DorsAttn_Post_47 | DorsAttn | | | | • | 6,-54,61 | 35.32 |
| 736 | 7Networks_RH_DorsAttn_Post_52 | DorsAttn | • | | | | 19,-48,70 | 22.36 |
| 739 | 7Networks_RH_DorsAttn_FEF_3 | DorsAttn | | • | | | 25,4,53 | 33.80 |
| 740 | 7Networks_RH_DorsAttn_FEF_4 | DorsAttn | • | | | | 34,-7,59 | 31.99 |
| 801 | 7Networks_RH_SalVentAttn_Med_10 | SalVentAttn | | | • | | 13,-39,45 | 35.70 |
| 804 | 7Networks_RH_SalVentAttn_Med_13 | SalVentAttn | • | | | | 9,-43,51 | 34.99 |
| 124 | 7Networks_LH_SomMot_43 | SomMot | | | | • | -52,-4,47 | 31.23 |
| 149 | 7Networks_LH_SomMot_68 | SomMot | | | | • | -3,-33,59 | 27.31 |
| 673 | 7Networks_RH_SomMot_92 | SomMot | • | | | | 20,-13,72 | 35.28 |
| 677 | 7Networks_RH_SomMot_96 | SomMot | | | | • | 13,-14,73 | 31.03 |
| 1004 | 7Networks_LH_SubCort_Hipp | SubCort | • | | | | -26,-24,-13 | 31.94 |

### Supplementary Table 3

| COMPLEXITY - EXPLORATORY DATASET | | | TIMESCALES | | | | | |
|---|---|---|---|---|---|---|---|---|
| ROI | Region Name | Network | 205 | 143 | 109 | 88 | LPI Coord (x,y,z) | sum(-logp) |
| 638 | 7Networks_RH_SomMot_57 | SomMot | | • | • | • | 9,-9,47 | 36.24 |
| 855 | 7Networks_RH_Cont_Par_13 | Cont | | | • | | 38,-68,50 | 27.95 |
| 861 | 7Networks_RH_Cont_Temp_2 | Cont | | | • | | 60,-39,-19 | 23.77 |
| 446 | 7Networks_LH_Default_PFC_31 | Default | | | • | | -21,49,31 | 26.39 |
| 689 | 7Networks_RH_DorsAttn_Post_5 | DorsAttn | | | • | | 59,-53,-16 | 33.37 |
